# The ultrastructure of subarachnoid trabeculae in non-human primates varies by tissue region within the central nervous system

**DOI:** 10.64898/2026.08.28.747903

**Authors:** Colin Riley, Olivia Mihalek, Rrita Daci, Lily Bigelow, Martha Fowler, David I. Sandberg, Christopher F. Janssen, Rachael W. Sirianni

**Affiliations:** Department of Neurological Surgery, UMass Chan Medical School, Worcester, MA, United States of America; University of Massachusetts Amherst, Department of Biomedical Engineering, Amherst, MA, United States of America; Vivian L. Smith Department of Neurosurgery, McGovern Medical School, University of Texas Health Science Center at Houston (UTHealth), Houston, TX, United States of America; Department of Bioengineering, Rice University, Houston, TX, United States of America; Department of Pediatric Surgery, McGovern Medical School, UTHealth, Houston, TX, United States of America; Center for Laboratory Animal Medicine and Care, McGovern Medical School, UTHealth, Houston, TX, United States of America; Department of Biochemistry and Molecular Biotechnology, UMass Chan Medical School, Worcester, MA, United States of America

**Author notes:** Corresponding Author Corresponding Author: Rachael W. Sirianni, PhD, Department of Biochemistry and Molecular Biotechnology University of Massachusetts Chan Medical School, 364 Plantation Street, Worcester, MA 01605. these authors contributed equally to this work (Colin Riley, Olivia Mihalek).

**Keywords:** Central nervous system, Subarachnoid trabeculae, Cerebrospinal fluid

## Abstract

The subarachnoid space (SAS) plays an important role in central nervous system (CNS) physiology, housing cerebrospinal fluid (CSF), providing nutrients, clearing waste, and protecting CNS tissues from injury. The SAS also contains an intricate arrangement of collagenous fibers, known as subarachnoid trabeculae, which serve as biomechanical support for the subarachnoid space and modulate the flow of CSF. Despite their myriad of roles in health and disease, the ultrastructure of trabeculae is incompletely described, with a specific gap in understanding how fiber structure varies across different regions of the CNS. In this work, we used optimized fixation techniques and scanning electron microscopy (SEM) imaging to study ultrastructural differences in trabeculae that were obtained from different regions of the nonhuman primate (NHP) brain and spinal cord. Qualitative assessments included evaluation of the relative prevalence of different kinds of fiber networks by region, leading us to a redefinition of terminology for fiber morphology. Quantitative measurements yielded dramatic differences in fiber diameter, density, and network porosity between different regions of the CNS. Broadly, trabeculae supporting cauda equina, cervical spinal cord, and exiting nerve roots were observed to have a very high density of small diameter fibers, whereas larger fibers and more complex arrangements were observed in other regions of the CNS. These data contribute to a growing field understanding of CNS-region specific differences in tissue ultrastructure.

**Graphical Abstract:** 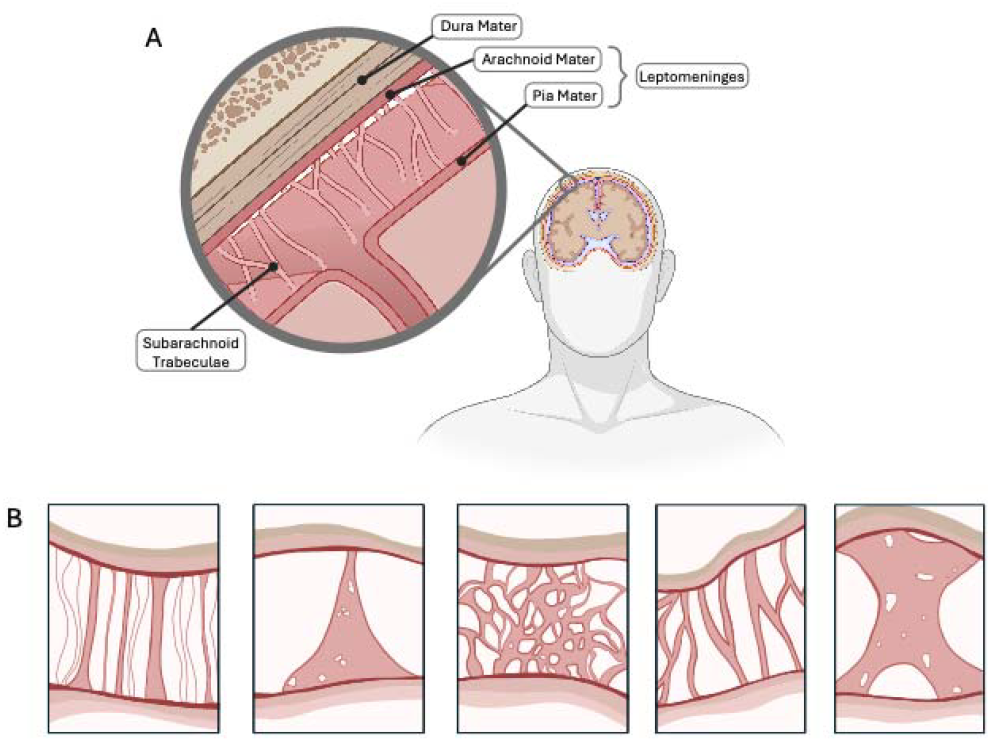

## 1. Introduction

The central nervous system (CNS) contains three layers of tissue that protect the brain and spinal cord. This includes dura, arachnoid, and pia mater (from superficial to deep),[1, 2] which, collectively, are referred to as the meninges. The space between the arachnoid and pial layers is known as the subarachnoid space (SAS), and it plays a critical role in providing mechanical support to the SAS, modulating flow of CSF, and mediating both health and disease in the CNS [3-5].

The dura mater is the outermost layer and contains two fibrous layers. The most superficial layer of the dura is called the periosteal layer, which lays adjacent to bone, followed by the deeper meningeal layer, which is in contact with the arachnoid mater. The periosteal layer of the dura is composed of fibroblasts, osteoblasts, and extensive amounts of extracellular collagen. It provides structural strength to the membrane and enables firm attachment to the skull’s sutural sites. The meningeal layer of the dura contains fibroblasts and a flattened mesothelial cell basement lining that maintains direct contact with arachnoid mater [6, 7]. Although the dura mater has often been viewed as a biologically inert aspect of CNS physiology, recent observations highlight an important role for dura in influencing CSF dynamics within the CNS [8].

The arachnoid mater is named for its spider-web-like projections that bridge the sulci [6, 7], although in reality these structures are likely more accurately described as cobwebs (possessing a messy, 3D structure) than as flat, 2D spiderwebs. Arachnoid mater is comprised of epithelial-like cells that facilitate CSF clearance to the venous system through arachnoid granulations [9]. Projections of collagenous fibers extend from the arachnoid mater to the pia mater, and these fibers are collectively called subarachnoid trabeculae (SAT)[6, 10]. Compared to the arachnoid layer, the pia mater is more complex and relatively elastic, consisting of reticular collagen and forming the epipial layer and the intima pia. This dual-layered structure allows the pia mater to form small channels adjacent to blood vessels throughout the brain and meninges. These channels facilitate the exchange of cellular and fluid materials with interstitial fluid of the parenchyma, eventually clearing to the cervical lymphatic system. The arachnoid and pia mater are collectively referred to as the leptomeninges, which encompass the SAS. In addition to the flow of CSF throughout the SAS, the leptomeninges form a sheath that surrounds vasculature, creating a space that is commonly referred to as the perivascular space (PVS), or, alternately, Virchow-Robin channels. The PVS is thought to play a critical role in facilitating fluid clearance from the CNS [6, 9]; we have also reported that the PVS also serves as a clearance route for intrathecally administered colloids [11].

SAT are primarily composed of type I collagen fibers, which are enveloped by resident fibroblast-like cells, termed leptomeningeal cells [1]. Mechanically, SAT provide viscoelastic support, dampening relative motion between the brain and skull and stabilizing neurovascular structures [2]. Additionally, SAT have been shown to influence CSF flow and pressure within the SAS, and this is an area of ongoing discussion [2]. Clinical pathologies linked to the leptomeninges and the microanatomical structures within the SAS include, but are not limited to, traumatic brain injury [12-16], hemorrhage [10, 15-19], and cancer metastasis [20-23]. Understanding the diverse morphologies and functions of SAT is essential for advancing our knowledge of both CNS biomechanics and disease states, ultimately improving clinical outcomes.

Despite the growing clinical relevance of the SAS and SAT for human health and disease, systematic or quantitative characterization of SAT remains limited [24]. Although the fiber composition is generally agreed on, the architecture of SAT is poorly defined. There remains debate regarding anatomical descriptions of subarachnoid pial connections and inconsistent use of SAT morphology nomenclature [25, 26]. Many reports on SAT ultrastructure focus on specific samples or regions, with less comparative analyses of how these fiber structures vary within the CNS. For example, SAT are often depicted as individual columns arranged in a radial pattern around the surface of the parenchyma. In fact, this depiction is primarily a result of extensive characterization of trabeculae within the optic nerve, which is relatively easy to access compared to other tissues of the CNS. Our published work and others make clear that the structure of SAT is considerably more complex and depends on the region of the CNS that is examined [6, 27].

Here, we sought to systematically examine differences in SAT ultrastructure of samples taken from multiple regions of the CNS. Our analysis includes both qualitative and quantitative assessment of fiber size, density, and morphology. We hypothesized that the ultrastructure of SAT varies based on anatomical location within the CNS. To test this hypothesis, we examined SAT collected from rhesus macaque non-human primates (NHPs), using scanning electron microscopy (SEM) to specifically study ultrastructure at the sub-micron level. By integrating quantitative and qualitative approaches, we aim to develop better, more comprehensive insight into the size and shape of these fibers to advance field-understanding of SAT ultrastructure.

## 2. Methods

### 2.1. Tissue Collection

All procedures were conducted in accordance with protocols approved by the University of Texas Health Science Center at Houston Animal Welfare Committee. The tissues utilized in this work were obtained from rhesus macaque NHPs (two males and two females, 4-7 years of age). These cadavers were previously used in a different study to evaluate the tolerability of intrathecal drug treatments (see prior publication for details)[28]. Once animals were deeply anesthetized with isoflurane, they were euthanized by exsanguination and transcardial perfusion with saline followed by 10% formalin fixative. Brain and spinal cord samples with the dura, arachnoid, and pia mater intact, were collected by craniotomy and laminectomy, respectively. Samples were collected from 12 regions of interest: the Sylvian Fissure, pituitary gland, thalamus, cerebellum, pineal gland, brainstem, olfactory nerve, optic nerve, cervical spinal cord, thoracic and lumbar spinal cord, cauda equina, and exiting nerve roots. After samples were collected, they were stored in 10% formalin at room temperature for 24-48 hours prior to processing.

### 2.2. Tissue Processing

Following tissue collection and initial fixation, samples were post-fixed in 3% glutaraldehyde solution at 4°C overnight, before being progressively dehydrated in increasing concentrations of ethanol (30%, 40%, 50%, 60%, 70%, 80%, 90%, and 100%). The samples soaked in each ethanol concentration for 10 minutes, until reaching the 100% concentration condition, where they soaked for two consecutive 10-minute periods. After dehydration, critical point drying was utilized to preserve the trabecular fiber structure and prepare the samples for imaging. The samples were then mounted with carbon adhesive on aluminum stubs and sputter coated with gold for 160 seconds at 40mAmps. Using a FEI Quanta 400 environmental SEM, samples were imaged with a secondary electron beam with a power of 20kV with a 10mm-17mm working distance, depending on image magnification.

### 2.3. Image Analysis

SAT images were analyzed using the Fiji ImageJ software (v2.3.0, NIH). Images were first converted to binary by adjusting the threshold value to maximize the contrast between fibers and background. Due to the intricacy of the NHP trabecular fibers, automated measurements were not accurate. Ultimately, a human experimenter manually measured fiber density and fiber diameter through systematic sampling. A minimum of three samples were measured for each of the 12 regions of interest. Fiber width was measured orthogonal to major fiber direction at a minimum of three locations per fiber. Individual fiber widths were recorded and then averaged to find the mean fiber diameter per tissue region of interest. To quantify fiber density, a 50µm^2^ box was drawn and individual fibers were counted within that box.

### 2.4. Data Analysis

One-way or two-way ANOVA followed by posthoc Tukey’s test was performed using GraphPad Prism 9.3.1 software. Statistical significance is reported at an alpha level of 0.05. Results are reported as mean +/-standard deviation for a minimum of three experimental replicates except where otherwise noted.

## 4. Results

The most recent major categorization of SAT morphologies includes rods, branches, trees, sheets, and networks **(Figure 1A)**[6, 8, 12]. Following our initial review of images, we determined that this terminology was not ideal for the distribution of morphologies found in this work. For example, the fibers that were previously called ‘trees’ and ‘branched’ shared many features in common, and, in our analysis, did not differ significantly in diameter, length, or density. Therefore, we combined these two morphologies into a single group termed ‘branched’ **(Figure 1B)**. We also identified two distinct forms of ‘networks’, one in which fibers are aligned and one in which fibers are multidirectional. We therefore divided ‘networks’ into ‘aligned networks’ and ‘veil-like networks’. This yielded five terms for classifying morphology of SAT: rods, branches, sheets, aligned networks, and veil-like networks.

**Figure 1:**
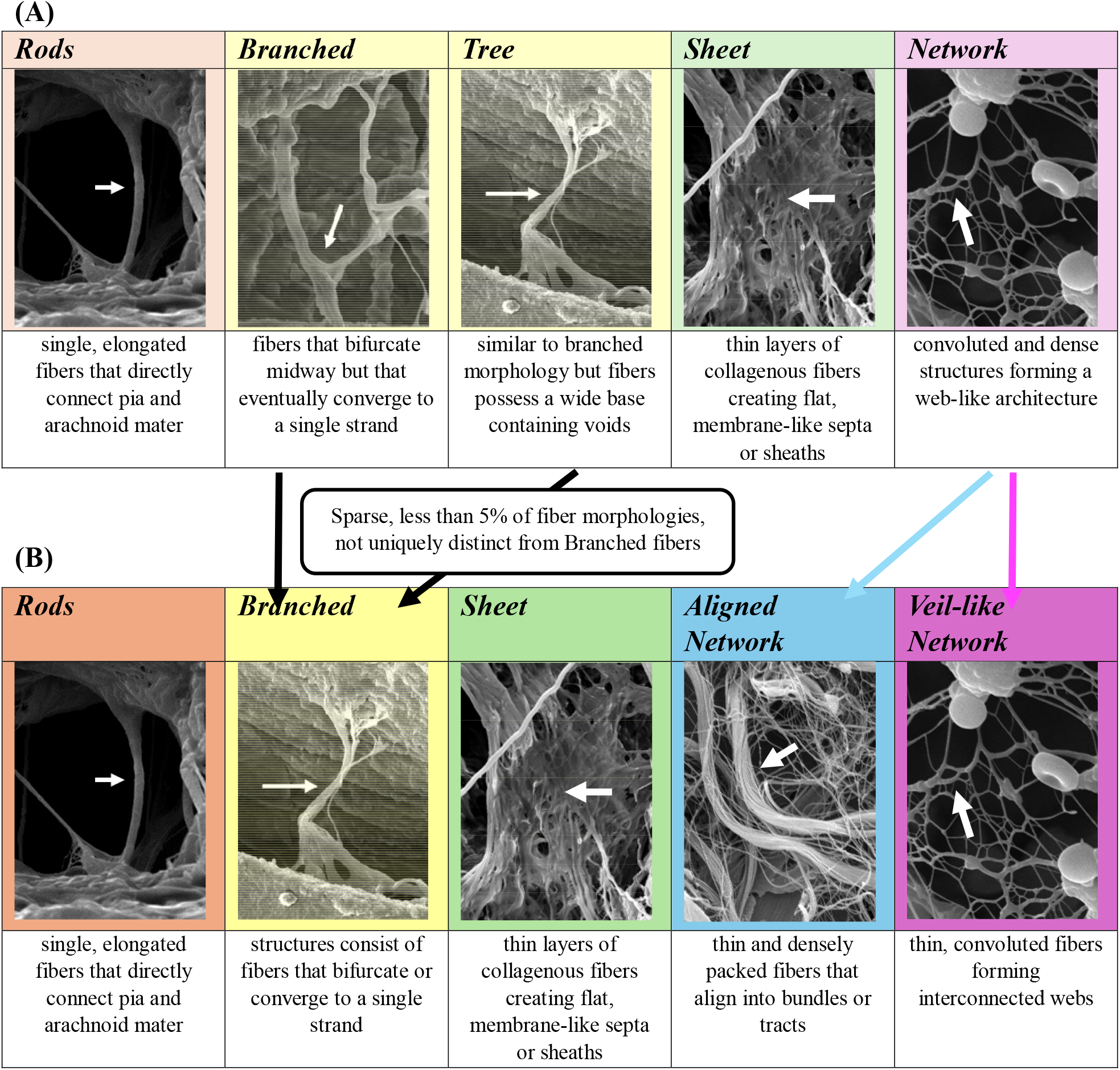
Proposed revision to typical descriptive terms for SAT. **(A)** SAT were previously categorized in five groups: rods, branches, trees, sheets, and networks. (B) We propose reclassifying these structures by combining ‘branches’ and ‘trees’ while also splitting ‘aligned networks’ and ‘veil-like networks’.

Mean fiber diameter was calculated by measuring hundreds of fibers in total across 12 CNS tissues **(Fig. 3A)**. Fiber diameters ranged across a factor of a thousandfold, from ∼90nm to ∼100um. In general, brain tissue and and cranial nerves exhibited larger fiber diameters compared to those observed in the spine. Notably, SAT examined in regions around the thalamus were characterized by significantly larger fiber diameters (p < 0.0001) compared to SAT in all other tissues. The cervical spinal cord, cauda equina, and exiting nerve roots displaying significantly smaller fiber diameters compared to all other groups (p < 0.0001). Group means, error, sample size, and statistical tests for fiber diameter are described in detail in **Sup. Table 1**.

**Figure 2:**
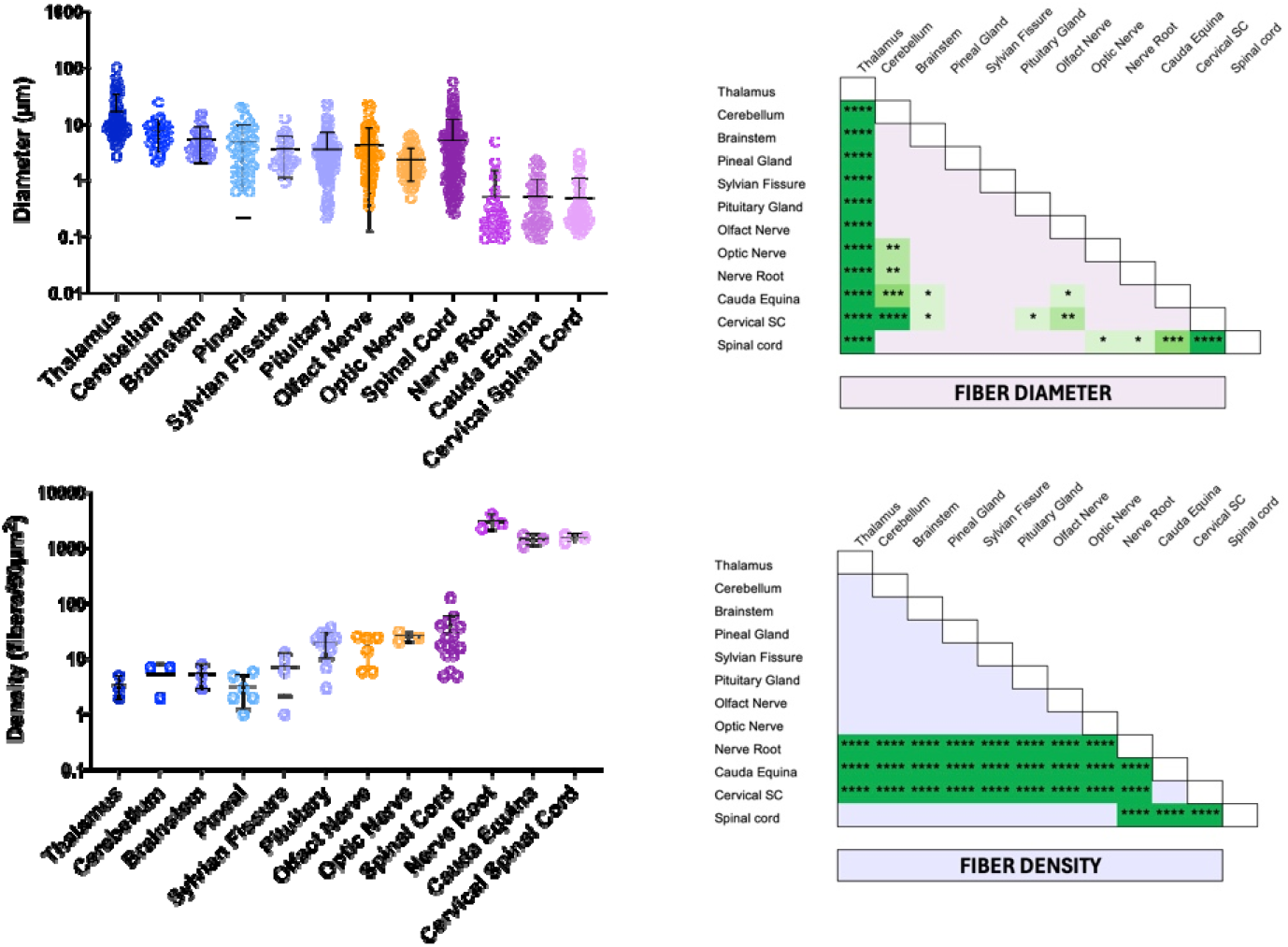
Trabeculae diameter and density were assessed on a per-fiber basis for 12 CNS tissue regions. Statistical significance is denoted by *p<0.05, **0.01, ***p<0.001, ****p<0.0001.

**Figure 3:**
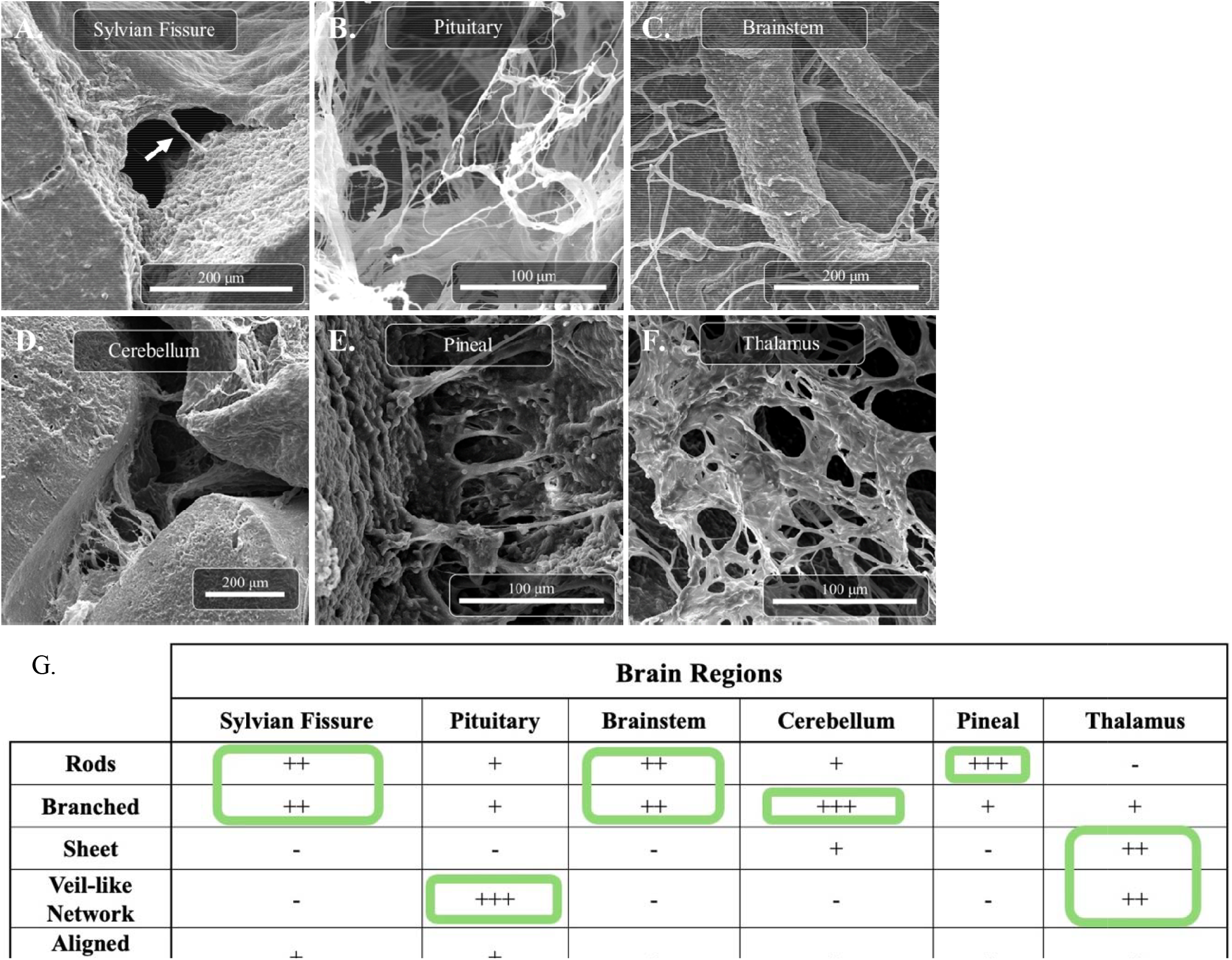
SEM images of SAT observed in different regions of the brain. Brain regions that were studied include **(A)** Sylvan Fissure, **(B)** Pituitary (Infundibulary Recess), **(C)** Brainstem, **(D)** Cerebellum, **(E)** Pineal Gland, and **(F)** Thalamus. Each of the types of SAT morphologies described in Figure 2 are evident in these panels: **(A)** rods, **(B)** veil-like networks, **(C-E)** rods and alined networks, **(F)** sheet. **(G)** Following analysis, relative morphologies were grouped based on the five categories described above and ranked based on prevalence per CNS region measured.

Fiber density was calculated by averaging the number of fibers present in 50µm^2^ regions of interest **(Figure 2B**. Densities were measured to be as low as 1 fibers/50µm^2^ (e.g., pineal gland and pituitary fissure) and up to 4075 fibers/50µm^2^ in the exiting nerve roots. The cervical spinal cord, cauda equina, and exiting nerve roots had significantly higher fiber densities (p < 0.0001) compared to all other CNS regions **(Sup. Table 2)**. These three regions also contained trabeculae with the smallest average diameters **(Sup. Table 3)**.

### 3.1. Brain samples – density and diameter

In general, fiber diameters assessed in samples collected from the brain were highly variable, ranging from 0.21µm in the pituitary region to 102µm in the thalamic region. Although the properties of individual fibers varied, we observed distinct patterns of distribution for different classification of fibers as a function of tissue region. The Sylvian Fissure separates the frontal and parietal lobes from the temporal lobe, and SAT in this region are observed to extend from the pia mater to arachnoid mater with predominately rod-like and branched SAT morphology **(Figure 3A, G)**. We note that the subarachnoid space associated with the Sylvian Fissure is relatively larger than other subarachnoid adjacent spaces. Both rod and branched SAT were observed with varying prevalence in tissues adjacent to the brainstem (mixed classification), cerebellum (predominately branched), and pineal gland (predominately rod-like) **(Figure 3C-E)**. We observed a relatively lower prevalence of rod-like and branched SAT in the pituitary region, as well as a low prevalence of rod-like SAT in the cerebellum and a lack of rodlike SAT in the thalamus. A substantial number of sheet SAT were identified in the thalamus (**Figure 3F**), and the pituitary region had a particularly high level of veil-like networks (**Figure 3E**). In some instances, sheets took on the appearance of nearly solid membranes possessing variably dispersed pores possessing diameters between 2µm and 30µm **(Figure 3F)**. Cells, including red blood cells, were frequently observed in these samples (examples shown in **Figure 3E)**, which are most likely an artifact of tissue collection.

Fiber density measurements were generally consistent between different regions of the brain, with the exception of the pituitary region. The average density in the pituitary samples was 19.8 fibers/50µm^2^, while the other five regions (thalamus, pituitary, brainstem, cerebellum, and pineal gland) had an average density of 4.9 fibers/50µm^2^. Interestingly, from the brain regions we measured, the pituitary gland had the highest average density, yet the lowest average diameter (average density = 19.8 fibers/50µm^2^, average diameter = 3.5µm). The opposite is seen in th thalamus, which had the lowest average density and the highest average diameter (average density = 3.3 fibers/50µm^2^, average diameter = 16.5µm). Furthermore, the pituitary region was the only brain region observed to have a high prevalence of networks, specifically the veil-like network **(Sup. Table 1)**.

### 3.2. Spinal Cord – Density and Diameter

We next examined SAT structures within cranial nerves, spinal cord, and exiting nerve roots. Fiber diameter in cranial nerves I and II (the olfactory and optic nerves, respectively) ranged from 0.35µm to 23µm in the olfactory nerve, while a fiber diameter range of 0.48µm to 6.6µm in the optic nerve. Qualitatively, the total span of the SAS surrounding the olfactory nerve tended to be larger than what was observed in the optic nerve **(Figure 4A-B)**. The predominant fiber orientation of these two regions included rod-like and branched fibers, although occasional aligned networks were observed. These fibers created narrows clefts and tracks within the otherwise spacious SAS.

**Figure 4:**
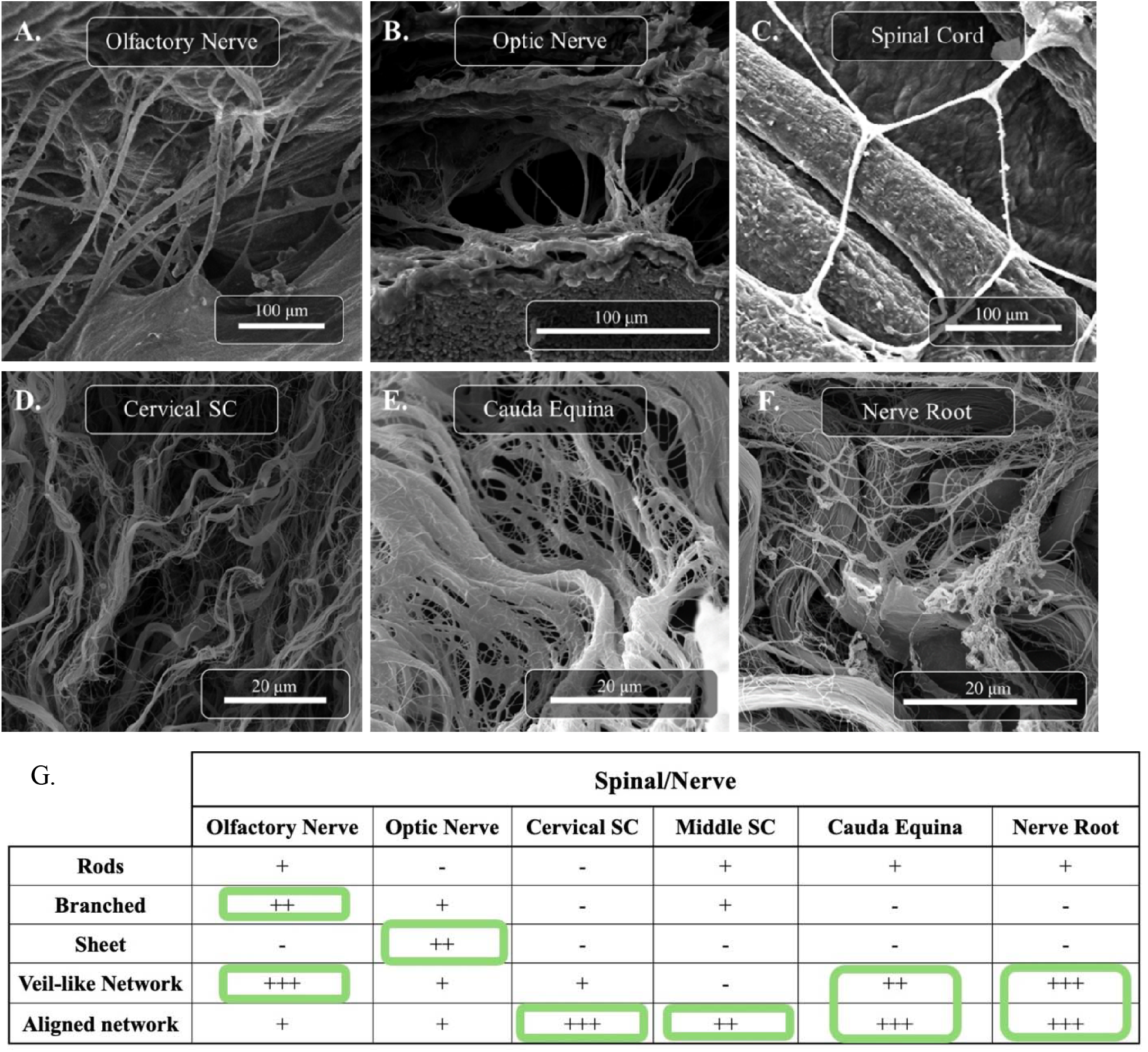
SEM images of SAT observed in different cranial nerves and regions of the spinal cord. Regions that were studied include (A) Olfactory Nerve, (B) Optic Nerve, (C) Spinal Cord, (D) Cervical Spinal Cord (SC), (E) Cauda Equina, and (F) Nerve Root. SAT tended to be columnar and branched in the cranial nerves. Within thoracic and lumbar spinal cord, cervical spinal cord, cauda equina, and nerve roots, fibers were observed to contain both aligned and veillike networks. **(G)** After analysis, fiber morphology was grouped into five distinct categories. Cranial nerves, specifically the olfactory and optic nerves, have more branched and columnar morphologies compared to spinal-cord regions, where network morphologies were more common.

Tissue samples from the spine were grouped by gross anatomy: thoracic and lumbar spinal cord, cervical spinal cord, cauda equina, and nerve roots **(Fig. 4C-F)**. Fiber diameters varied from as small as 0.092 µm (in the cauda equina) to as great as 57µm (in the spinal cord). These tissue regions were frequently supported by complex SAT ultrastructure, typically with multiple fibers connecting at vertices to form irregular polygons and veil-like networks. Although the individual fibers were frequently uniform in size and shape, when aggregated into tracts, these complex fibers yielded veil-like networks with many particularly small fibers that were packed together to create an effectively larger fiber. We note that many conventional imaging techniques do not possess the spatial resolution needed to identify sub-100nm fibers; in this work, SEM reveals the very small ultrastructure of individual fibers that are contained within larger bundles.

Compared to what was observed in the brain, trabeculae around the cervical spine, cauda equina, and nerve roots were far denser. The average density in these three regions were 1,778 fibers/50 µm^2^, 1,441 fibers/50 µm^2^, and 3,050 fibers/50 µm^2^, respectively. These regions possessed uniquely dense, aligned networks **(Fig. 4D-F)**. Many of the fibers in these regions were not only much smaller than fibers we observed elsewhere, but many were flat, with tracts resembling ribbons. We cannot exclude the possibility that there were smaller fibers contained within these ribbon-like structures, but, if present, these smaller fibers were not visible on these samples imaged by SEM. The cauda equina and nerve roots also contained veil-like networks, which contributed to the overall fiber density in the area. In summary, the fibers observed in the spinal cord and exiting nerve roots were generally smaller in diameter and more densely packed, generating both aligned and veil-like networks, whereas fibers within cranial nerves I and II displayed primarily columnar and branch structures **(Sup. Table 2)**.

## 4. Discussion

Growing scientific interest in the structure of SAT stems from the evolving literature suggesting that the subarachnoid space is not simply a static compartment, but a structurally complex, dynamically regulated microenvironment that plays a significant role in fluid transport, immunity, and disease pathology of the CNS. Computational fluid mechanical approaches have found that trabeculae structures significantly increase cerebrospinal fluid flow resistance, with some simulations reporting a 2.5-fold increase in pressure drop compared to models without trabeculae. Additionally, it has been found that trabeculae enhance micromixing by inducing vortices and recirculation zones within the SAS. SAT thereby directly disrupt laminar flow, increasing the complexity of CSF dynamics, and highlighting a point of clinical significance when related to drug biodistribution in the CNS [29]. Beyond the role in fluid dynamics, SAT provides mechanical support and viscoelastic cushioning, dampening the motion of the brain relative to the skull and increasing stability [2]. Significantly, there is emerging evidence that the presence and structure of SAT play a role in mediating disease progression [30]. Work by our group and others suggests that SAT ultrastructure could be iplicated in cancer metastasis in the CNS [17, 31].

The composition of SAT has been studied in both animal models and from human tissue samples. SAT are composed predominately of type 1 collagen fibers as confirmed by immunohistochemistry, proteomic analysis, and electron microscopy [12]. Immunohistochemical analysis of human optic nerve sections showed strong COL1A1 staining in pia and arachnoid mater. Recent studies have identified thin layers of fibroblast-like cells enveloping SAT and spanning between the pia and arachnoid membranes. COL1A1 mRNA was detected in cultured primary human meningothelial cells, confirming collagen as a primary structural component of SAT [22, 32]. Transcriptomic and morphological analysis have found that fibroblasts form a continuous lattice along the surface of SAT, suggesting they may play a role in maintaining structural integrity while also influencing cellular dynamics and signaling within the SAS [33-36]. Imaging and tracer studies have also illustrated that SAT contribute to the arachnoid membranes and septa that form cisterns and perivascular spaces within the SAS, influencing CSF flow and pressure dynamics [21, 36, 37]. Additionally, the variability of both the density and morphology of SAT can influence pressure/flow of CSF and may contribute to asymmetric clinical findings such as papilledema [21, 29].

Although some progress has been made to understand both composition and ultrastructure of SAT, current literature utilizes disparate terminology and lacks consensus on trabecular morphology and regional variability in architecture [8]. This is largely attributed to the diverse imaging techniques used to study SAT, all of which have unique advantages and disadvantages. MRI allows for a non-invasive, *in vivo* view of the entire SAS and some of the microstructures within it. While MRI is useful for assessing CSF flow and mapping gross anatomical features (e.g., vasculature) in the brain, its limited spatial resolution prevents accurate visualization of SAT ultrastructure. As a result, MRI can lead to underestimations in abundance and distribution of SAT within the SAS [8, 37]. In contrast, optical coherence tomography (OCT) provides high resolution three-dimensional imaging on the micrometer scale, which has yielded significant, novel observations regarding SAT ultrastructure on micrometer scale resolution. This technique has been usful, but limited tissue penetrance and required transvascular access restrict imaging to regions immediately adjacent to vasculature. Nevertheless, volume fraction analysis of OCT imaging has shown regional variability of subarachnoid trabeculae [1, 25, 38]. Scanning and transmission electron microscopy (SEM/TEM) provide nanometer-scale resolution, enabling detailed visualization of complex SAT architectures while sacrificing efficiency and throughput. The high-resolution images offer clear points of reference for the characterization of the various SAT morphologies. A majority of prior work with SEM have focused on individual regions of interest, such as the optic nerve. In this work, we focused on using SEM in a broader fashion to enable ultrastructural characterization across different regions of the CNS.

Our study provides quantitative and comprehensive assessment of SAT ultrastructure, and we identified clusters of trabeculae representing different aspects of the density:fiber diameter spectrum (results summarized in **Figure 6**). We observed that SAT in the brain and cranial nerves generally exhibit larger diameters than trabeculae associated with the spinal cord regions. Brain and cranial nerve trabeculae primarily consist of rod and branched-like morphologies, while the spinal cord regions exhibited a greater prevalence of complex trabeculae networks. Additionally, these regions tend to have less dense SAT structures (fiber density in brain regions: 3.2-25.6 fibers per 50 µm^2^). The structural characteristics of these larger fibers within the brain might suggest that they provide increased mechanical support for the larger CNS structures (fiber diameters in brain regions: 3.5µm to 16.5µm). For instance, research into traumatic brain injury has found that mechanical properties of the pia-arachnoid complex play an important role during impact or inertial loading. Studies have shown that regions of the CNS containing larger and higher density of fibers which directly influence tissue level mechanical properties by taking a protective role via dissipating mechanical loads and protecting the brain from injury [39-41]. Thus, the variation in SAT diameter across brain regions could potentially play a role in region-specific pathophysiology that results from injury.

**Figure 6:**
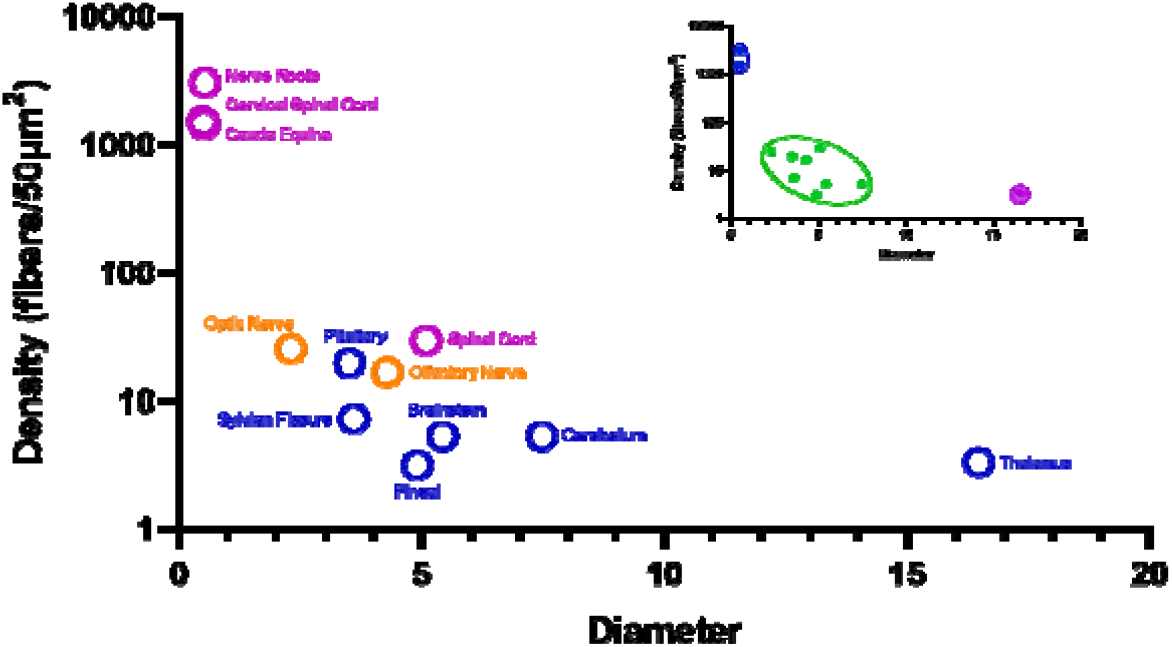
Relationship between average trabeculae diameter and density in each brain region of interest.

In contrast to the brain, SAT in the spinal cord, cauda equina regions, and exiting nerve roots are characterized by smaller fiber diameters (e.g., in the cauda equina: 91nm to 2.3µm). These regions typically display higher fiber densities (fiber density in spinal cord regions: 1441.7 fibers per 50 µm^2^). The transition from brain to the spinal cord yields a shift in SAT morphologies from large sparse rods and branched trabeculae to small and more convoluted, aligned and veil-like networks. Both the fiber diameter and fiber density change by nearly a factor of 10 when comparing brain trabeculae to trabeculae present in different regions of the spinal cord and exiting nerve roots.

As knowledge has grown regarding the SAS, so too have the terms used to classify and understand SAT ultrastructure morphology. SAT are often schematically depicted as individual columns radiating from the pia mater and extending completely to the arachnoid mater. Our work **(Fig. 2, 3**,**4)** and others emphasizes that this morphology is observed in some regions of the CNS, however, its prevalence is likely overstated in literature due to an overrepresentation of trabeculae from the optic nerve [6]. This tendancy to overrepresent trabeculae as singular, radial columns is likely due to the ease of access to the optic nerve relative to the difficulty of accessing and preserving SAT structures in the rest of the CNS. Our data (**Fig. 2, 3, 4**) emphasize that these radial columns are only a subset of what is in fact a highly diverse and complex arrangement of trabecular fibers that, in many instance, are so densely packed as to form sheets. While SAT terminology initialy focused on columns, pillars, and septa, growing investigation has expanded the use of terminology to include rod-like structures, branched forms, tree-like configurations, plate-like shapes, complex networks, branching strands, fenestrated sheets, and complex networks [1, 2, 6, 8, 21, 25].

To address this field-level complexity and deepen the understanding of ultrastrucrual features of SAT at the nanoscale level, we used SEM to examine tissues across 12 anatomically distinct regions of the CNS. Increasing the spatial sampling while utilizing a single imaging modality enabled us to compare morphologies and their regional prevalence, allowing us to form a revised classification system. Our schema consolidates overlapping categories (combining previous ‘tree-shape’ and ‘branched’ morphologies) and subdivides the broad network classification into distinct morphologies resulting in five SAT types: rods, branches, sheets, aligned networks, and veil-like networks. This approach to terminology in effect merged trabeculae that had been previously called “trees” into the category of “branches”, which is a relatively minor change. In our analysis, two groups emerged based on orientation of fibers. One subset of fibers were aligned in ribbon like tracts while the other displayed multidirectional fibers forming webs, thus leading to the suggested terms “aligned networks” and “veil-like networks, which more accurately describe the diversity in trabeculae ultrastructure that would be expected to impact biomechanical and physiological functions.

## Supplemental Figures (Tables)

**Supplemental Table 1:**
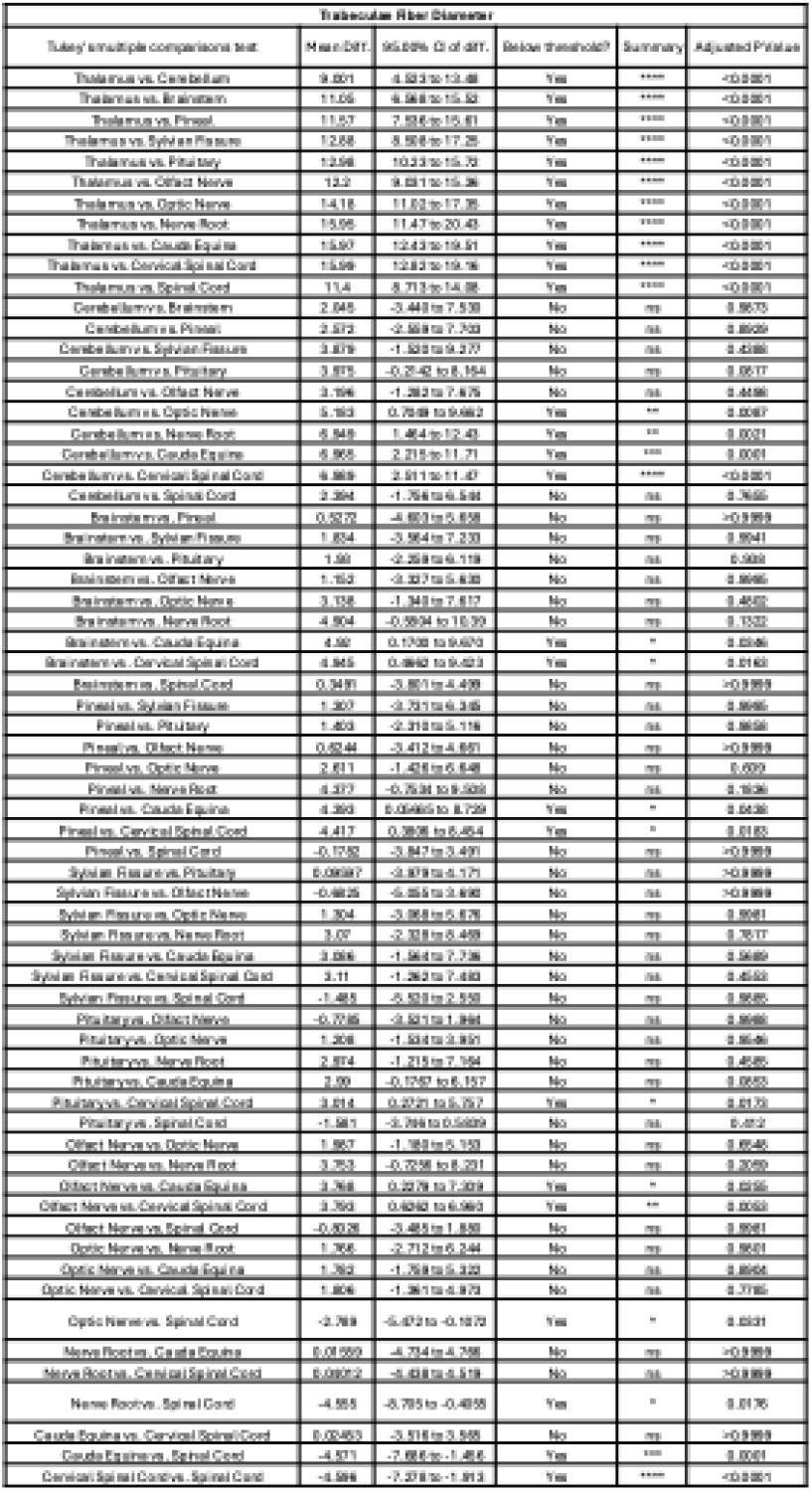
Tukey’s multiple comparison test results on trabeculae fiber diameter.

**Supplemental Table 2:**
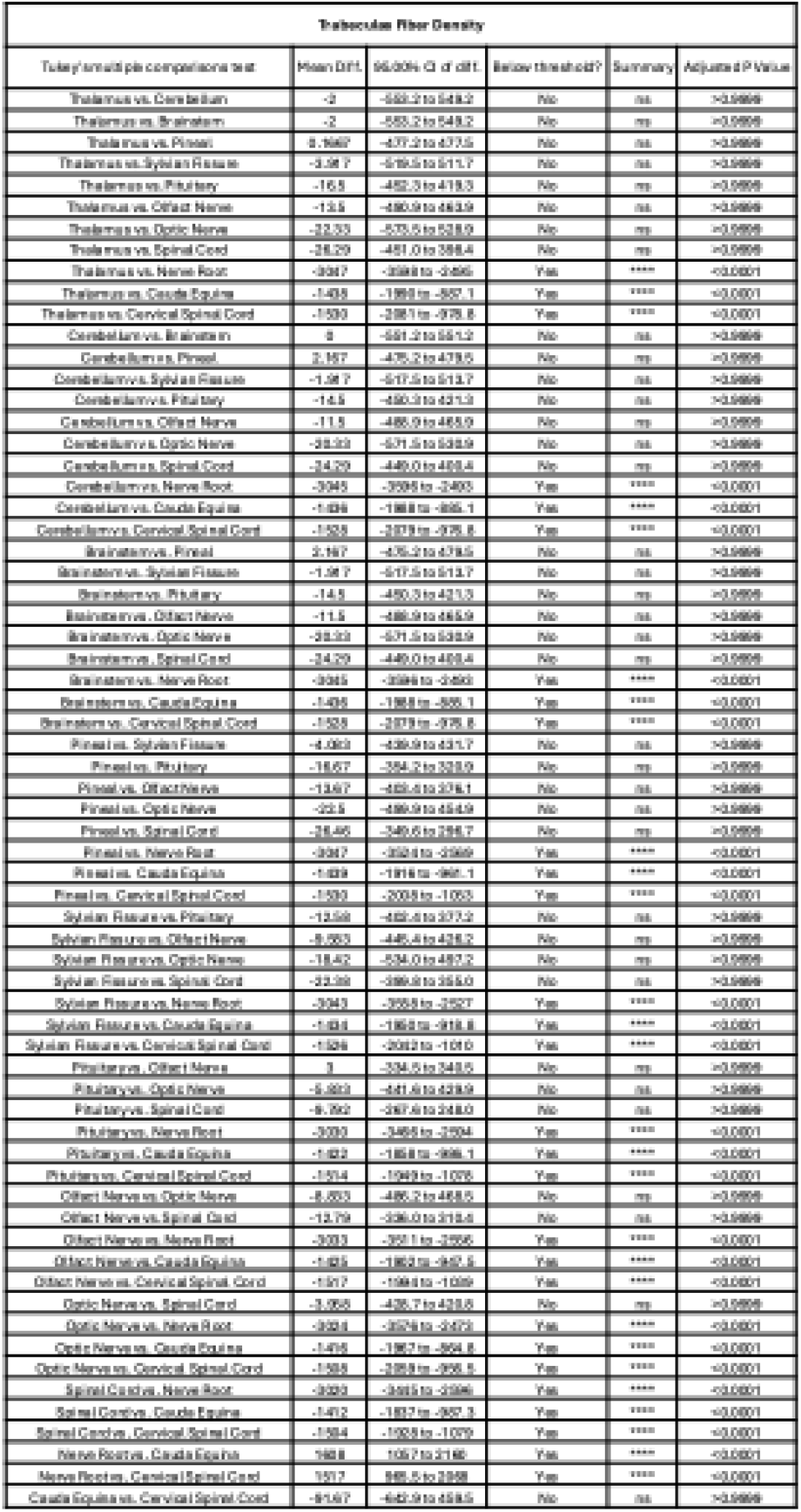
Tukey’s multiple comparison test results on trabeculae fiber diameter.

**Supplemental table 3:**
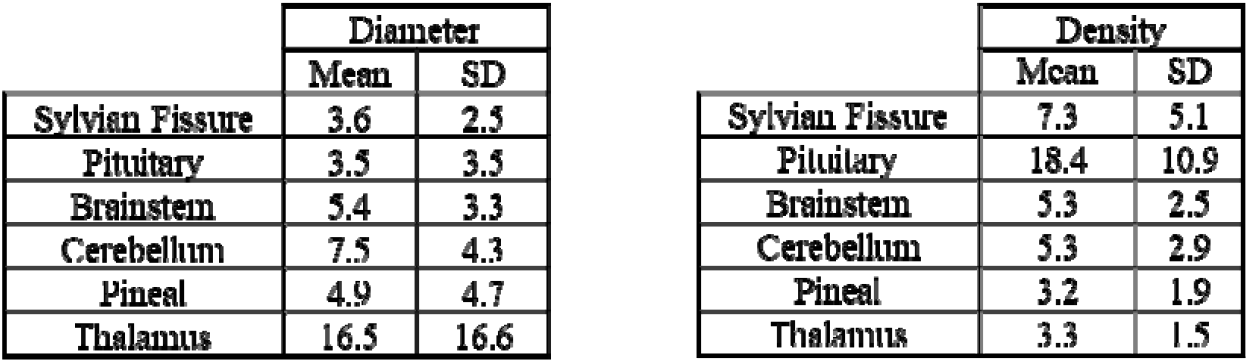
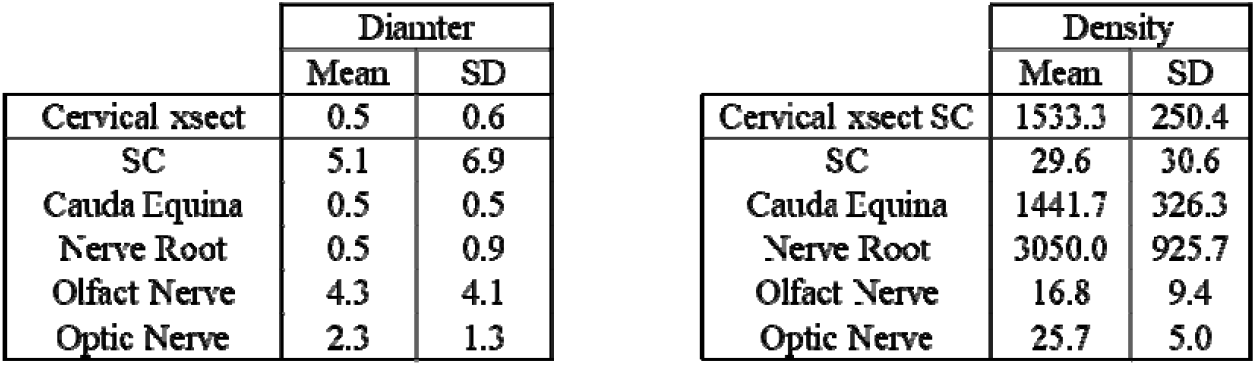
The average diameters and densities of trabeculae fibers by CNS region.

## Notes

### Competing Interest Statement

The authors have declared no competing interest.

